# Revealing the Physiological Origin of Event-Related Potentials using Electrocorticography in Humans

**DOI:** 10.1101/2021.02.12.430921

**Authors:** Hohyun Cho, Gerwin Schalk, Markus Adamek, Ladan Moheimanian, William G. Coon, Sung Chan Jun, Jonathan R. Wolpaw, Peter Brunner

## Abstract

The scientific and clinical value of event-related potentials (ERPs) depends on understanding the contributions to them of three possible mechanisms: (1) additivity of time-locked voltage changes; (2) phase resetting of ongoing oscillations; (3) asymmetrical oscillatory activity. Their relative contributions are currently uncertain. This study uses analysis of human electrocorticographic activity to quantify the origins of movement-related potentials (MRPs) and auditory evoked potentials (AEPs). The results show that MRPs are generated primarily by endogenous additivity (88%). In contrast, P1 and N1 components of AEPs are generated almost entirely by exogenous phase reset (93%). Oscillatory asymmetry contributes very little. By clarifying ERP mechanisms, these results enable creation of ERP models; and they enhance the value of ERPs for understanding the genesis of normal and abnormal auditory or sensorimotor behaviors.

## Introduction

The brain produces electrical responses to sensory, cognitive, and motor events. These responses, called event-related potentials (ERPs), can be detected by averaging the electrical activity recorded from electrodes placed on the scalp (electroencephalography (EEG)) (***Makeig et al., 2004***). ERPs have been used for decades to study different aspects of information processing in the brain (such as attention (***Coull, 1998***) or cognitive workload (***Isreal et al., 1980***)) or to diagnose specific neurological disorders (such as deficits in the auditory or visual system (***Coats, 1978***; ***Parisi et al., 2001***)).

ERPs are characterized by the type of event that causes them, e.g., auditory stimulation (resulting in auditory evoked potentials (AEPs)) or movements (resulting in movement-related potentials (MRPs)), and by the polarity (positive/negative) and latency (e.g., 100 ms) of the dominant peak in the response elicited by the event (***Picton et al., 1974***). Prior studies have hypothesized that three principal mechanisms give rise to ERPs (***Nikulin et al., 2007***): 1) additivity of event-related signal components (***Dawson, 1947***; ***Shah et al., 2004***; ***Mäkinen et al., 2005***; ***Turi et al., 2012***); 2) phase reset of underlying low-frequency (<40Hz) oscillatory activity (***Savers et al., 1974***; ***Makeig et al., 2002***; ***Hanslmayr et al., 2006***); and 3) asymmetry in oscillatory activity (***Nikulin et al., 2007***; ***Mazaheri and Jensen, 2008***; ***van Dijk et al., 2010***; ***Schalk, 2015***; ***Schalk et al., 2017***).

Together, these three mechanisms could account for much of the interactions between stimulus, neural oscillations, and ERPs. For the additivity mechanism, the stimulus may induce a direct response. For the phase reset mechanism, the stimulus may induce a phase reset in an ongoing oscillation. For the asymmetry mechanism, the stimulus may induce a variation in the biased oscillation (i.e., an oscillation with asymmetrically distributed peak/trough amplitudes). The differential effects of these three mechanisms produce an ERP. Consequently, while the additive mechanism directly affects the ERP, phase reset and asymmetry mechanisms indirectly affect the ERP through modulation of the ongoing oscillation (see ***Figure 1***, ***Figure 1–Figure Supplement 1***, and ***Figure 1–Figure Supplement 2*** for further details).

**Figure 1.**
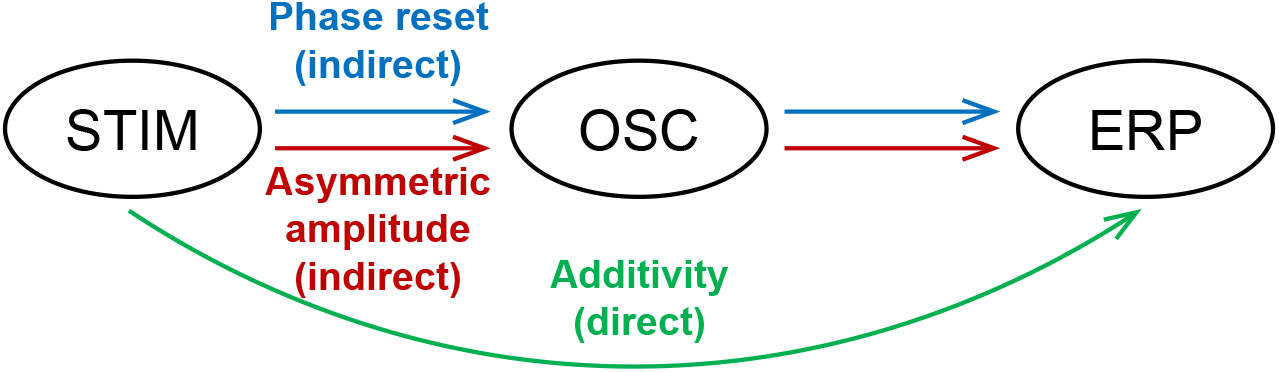
Generating mechanisms of ERPs. External stimuli (STIM) can give rise to ERPs directly through additivity, or indirectly through phase reset or asymmetric amplitude modulations of the ongoing oscillations (OSC). **Figure 1–Figure supplement 1.** Additivity and phase reset mechanism **Figure 1–Figure supplement 2.** Additivity and asymmetric amplitude mechanism

While these potential mechanisms have been described for decades (see ***Appendix 1***), there is still considerable debate about them and their unique contributions, particularly in the context of auditory evoked potentials (AEPs) (***da Silva, 2006***) and movement-related potentials (MRPs) (***Sauseng et al., 2007***). The principal challenge is that scalp-recorded EEG reflects a mixture of differ ent physiological phenomena that are produced by different areas of the brain, and decomposing EEG into its constituent components can produce different solutions (***Darvas et al., 2004***).

Moreover, determining the contribution of each of the three generating mechanisms to the gen eration of ERPs depends on the ability to completely decompose the neural signal into its additive, phase reset, and asymmetry components. Prior work has attempted to perform this decomposition only for additive contributions, and asymmetry in oscillation has only recently been proposed. In addition, investigating the role of phase reset in the generation of ERPs is diffcult. It requires that three preconditions are fulfilled (***Sauseng et al., 2007***): 1) the frequency band of the ongoing oscillation needs to be defined; 2) the frequency band of the ongoing oscillation must match that of the ERP component; and 3) the physiological sources of the ERP component and the ongoing oscillation need to match. Fulfilling these three preconditions requires access to spatially uncon-founded signals, which are only affected by local neural activity.

These requirements effectively limit the investigation of the contribution of the three generating mechanisms to signals recorded directly from the surface of the brain. To date, only one study has quantified the role of additivity in the generation of ERPs (***Turi et al., 2012***). In this study, Turi et al. investigated the generation of ERPs in monkeys using local field potentials and generalized their results to human subjects using MEG. However, this generalization depended on an independent component analysis (ICA), which assumes that the components of the signal are independent (***Hyvärinen et al., 2001***). This assumption conflicts with the precondition of phase reset, i.e., that ERPs can only be generated in the presence of an ongoing oscillation. Further, this study only considered the variance in the ERP amplitude without considering the relationship between the ongoing oscillation and the resulting ERP. These issues render the conclusions of Hyvärinen’s study to be less than certain.

Attempts to decompose EEG into its constituent components have resulted in uncertain (***Sauseng et al., 2007*; *Min et al., 2007*; *Fell et al., 2004***) or overtly conflicting (***Sauseng et al., 2007*; *Hanslmayr et al., 2006*; *Mazaheri and Jensen, 2006***) conclusions. This lack of clarity about the physiological origin of scalp-recorded evoked potentials greatly limits the detailed and accurate physiological interpretation of the results of thousands of studies using scalp-recorded ERPs, and the generation of more generalized models of brain function that such interpretation could inform.

In our study, we recorded signals from the surface of the brain to determine the physiological origin of auditory and movement-related potentials. We recorded electrocorticographic (ECoG) activity from eight human subjects while they executed a simple reaction-time task. In this task, the subjects responded to a salient auditory stimulus by pressing a push-button with the thumb contralateral to their ECoG implant. The subjects performed between 134 and 580 trials; their reaction time (277±110 ms) was comparable to that of similar previous studies (***Molholm et al., 2002***); and they responded within 1 s of the stimulus onset in 99.8±0.3% of all trials.

First, we determined those electrode locations at which ECoG high gamma activity (70–170 Hz), a widely accepted index of population-level cortical activity (***Edwards et al., 2005***; ***Darvas et al., 2010***; ***Ray and Maunsell, 2011***; ***Jenison et al., 2015***; ***Schalk et al., 2017***), increased in response to the auditory stimulus or the button press. Across all subjects, this procedure resulted in 22 and 15 task-related locations, respectively (***Figure 2***A and ***Figure 2–Figure Supplement 1***). We then determined the grand average MRP (***Figure 2***B) and AEP (***Figure 2***C) and verified their consistency across individual subjects (***Figure 2–Figure Supplement 2***). We further compared our results to those obtained from EEG in control subjects that performed the same experiment (***Figure 2–Figure Supplement 3*** & ***Figure 2–Figure Supplement 4***), and to those of similar experiments reported in the literature (***Figure 2–Figure Supplement 5***). To determine the specific contribution of each of the three possible generating mechanisms to the AEP and MRP responses, we assessed the fraction of the overall signal accounted for by each generating mechanism. Prior to this assessment, we verified that our data fulfilled the three preconditions for investigating phase reset (***Figure 3***). For this, we first determined that the frequency band of the ongoing oscillation across all eight subjects was within the 3–40 Hz band (***Figure 3–Figure Supplement 1***). Second, we verified that the N1 and P1 components were within the same band of the ongoing oscillation (***Figure 3***E). We fulfilled the third precondition by using ECoG signals for which the physiological sources of ERP components and ongoing oscillations are identical.

**Figure 2.**
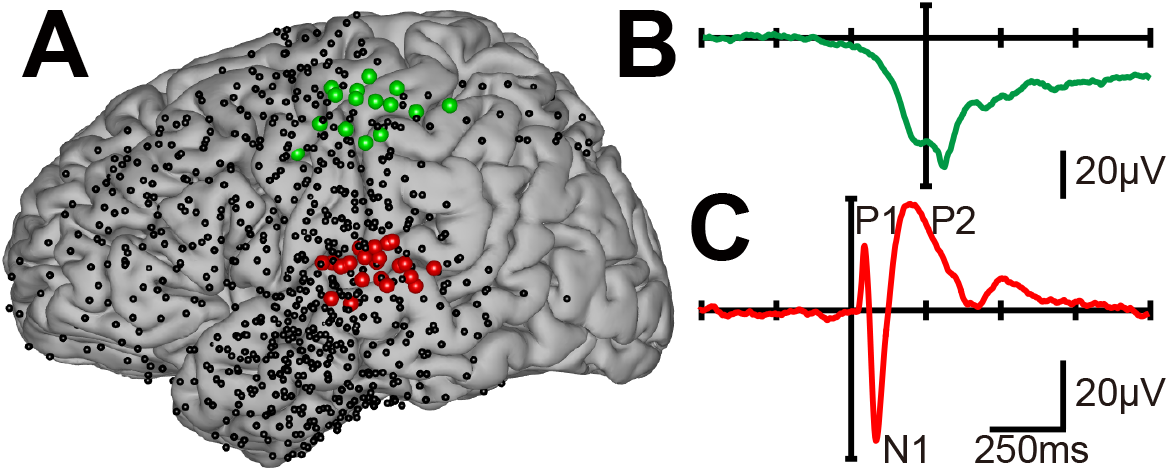
Location and shape of cortical ERPs. **A.** Electrode locations for all subjects (dots). Locations that exhibited task-related activity during auditory stimulation or motor movements are highlighted in red or green, respectively. **B.** The green trace shows the ERP produced by a button press, averaged across all task-related locations and subjects. **C.** The red trace shows the ERP produced by auditory stimulation, averaged across all task-related locations and subjects. These average ECoG ERPs are similar to ERPs reported in this or previous studies that used scalp-recorded EEG (***Figure 2–Figure Supplement 5***). **Figure 2–Figure supplement 1.** Task-related locations for each ECoG subject. **Figure 2–Figure supplement 2.** AEPs and MRPs for each ECoG subject. **Figure 2–Figure supplement 3.** Grand average AEP and MRP across all EEG subjects. **Figure 2–Figure supplement 4.** AEPs and MRPs from EEG for each EEG subject. **Figure 2–Figure supplement 5.** ERPs comparisons with EEG literature.

**Figure 3.**
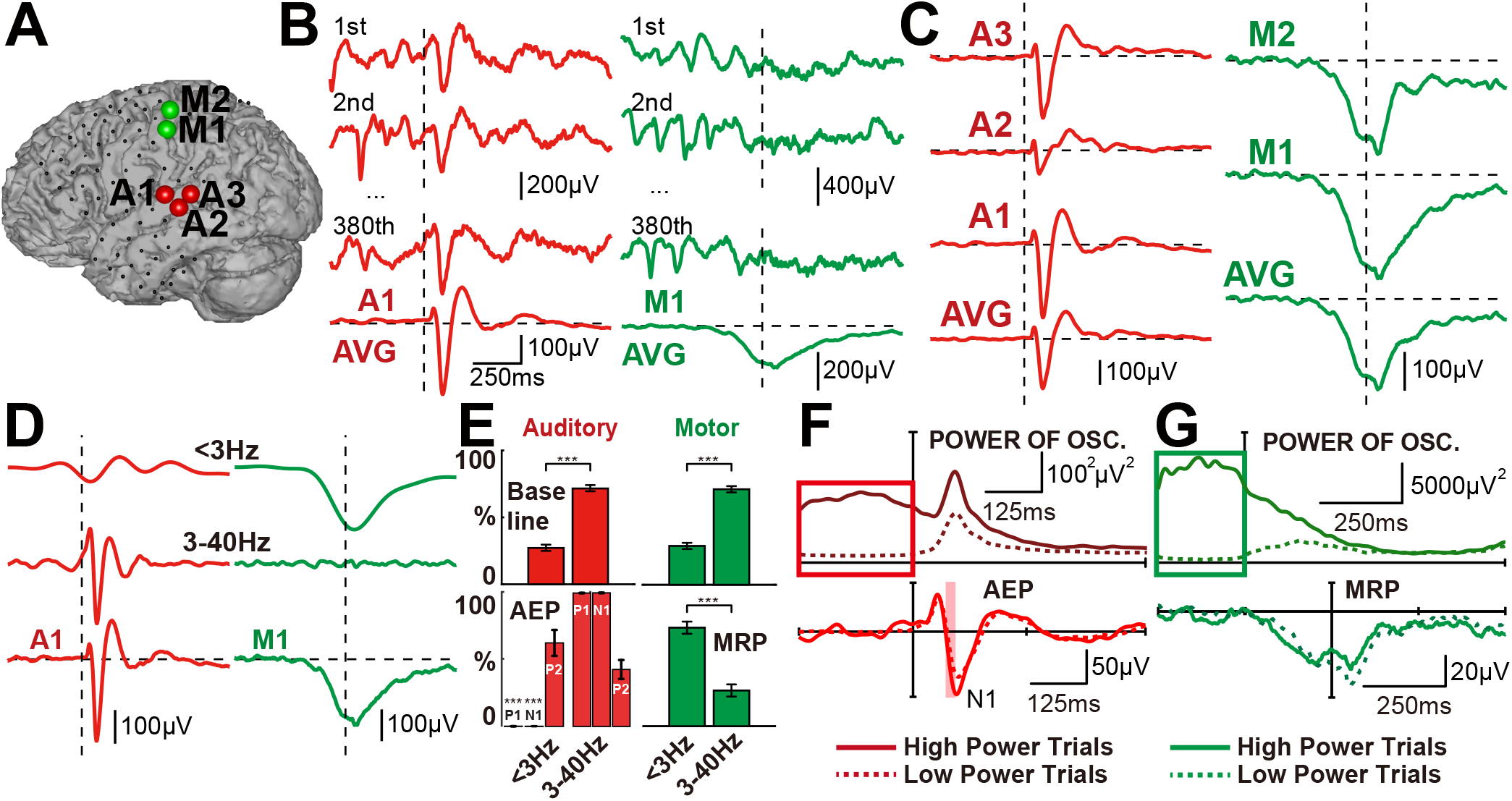
Overview of ERP analyses and their results. **A.** Locations exhibiting evoked potentials resulting from auditory stimulation (red dots) or a button press (green dots) in subject S3. **B.** Time course of ECoG activity during auditory stimulation (left) and button presses (right) for locations A1 and M1, and their across-trial average. Single-trial ECoG responses at location A1 are phase-locked at stimulus onset, and demonstrate the same N1, P1, and P2 components as seen in the across-trial average. In contrast, single-trial ECoG responses at location M1 are not phase-locked at movement onset, and thus no evoked potentials are exhibited in the average across all trials. Instead, a slow cortical potential arises from the average across all trials. **C.** Average AEPs (red traces on the left) and MRPs (green traces on the right) for locations A1-3 and M1-2 and their average in subject S3. All auditory locations exhibit a clear N1, P1, and P2 components, and all motor locations exhibit a prominent slow cortical potential. **D.** Time courses of ERPs from locations A1 and M1 in subject S3 in two different frequency bands (<3 Hz and 3–40 Hz). The characteristic components of the AEP are captured by the 3–40 Hz band. In contrast, the slow negative potential in the MRP can only be seen in the <3 Hz band. **E.** ECoG power in the <3 Hz and 3–40 Hz bands for baseline (−400 to 0 ms) and ERP (0 to 400 ms) periods (top and bottom, respectively), calculated across all task-related locations and all subjects. Baseline activity is mostly comprised of 3–40 Hz band power (p<0.001, paired t-test). The P1 and N1 components of AEPs are comprised of 3–40 Hz band power (p<0.001, paired t-test), while MRPs are mainly comprised of <3 Hz band power (p<0.001, paired t-test). **F.** Power (top) and shape of AEPs (bottom) in the 3–40 Hz band for trials with the highest (solid) and lowest (dashed) 10th percentile of pre-stimulus power (calculated per task-related location, averaged across all locations and subjects). Higher pre-stimulus power results in higher N1 amplitudes in AEPs (p<0.05, t-test, FDR corrected for N=22). **G.** Power (top) and shape of MRPs (bottom). Pre-stimulus power does not markedly affect the shape of MRPs (p<0.05, t-test, FDR corrected for N=15). **Figure 3–Figure supplement 1.** Determining the frequency band of the ongoing oscillation. **Figure 3–Figure supplement 2.** Relationship between ongoing oscillatory power and reaction time.

## Results

After verifying that all three preconditions for our phase reset analysis were fulfilled, we determined the contribution of each of the three generating mechanisms by removing their influence from the individual trials. For the additivity mechanism, we removed frequency components out-side of the ongoing oscillation. For the phase reset mechanism, we recomposed each trial’s signal within the frequency band of the ongoing oscillation with constant phase velocity (***Freeman, 2004***). For the asymmetric amplitude mechanism, we subtracted the asymmetric bias from each trial’s signal within the frequency band of the ongoing oscillation (***Nikulin et al., 2007***; ***Mazaheri and Jensen, 2008***; ***Schalk, 2015***). ***Figure 4*** illustrates this process for each of the three generating mechanisms.

**Figure 4.**
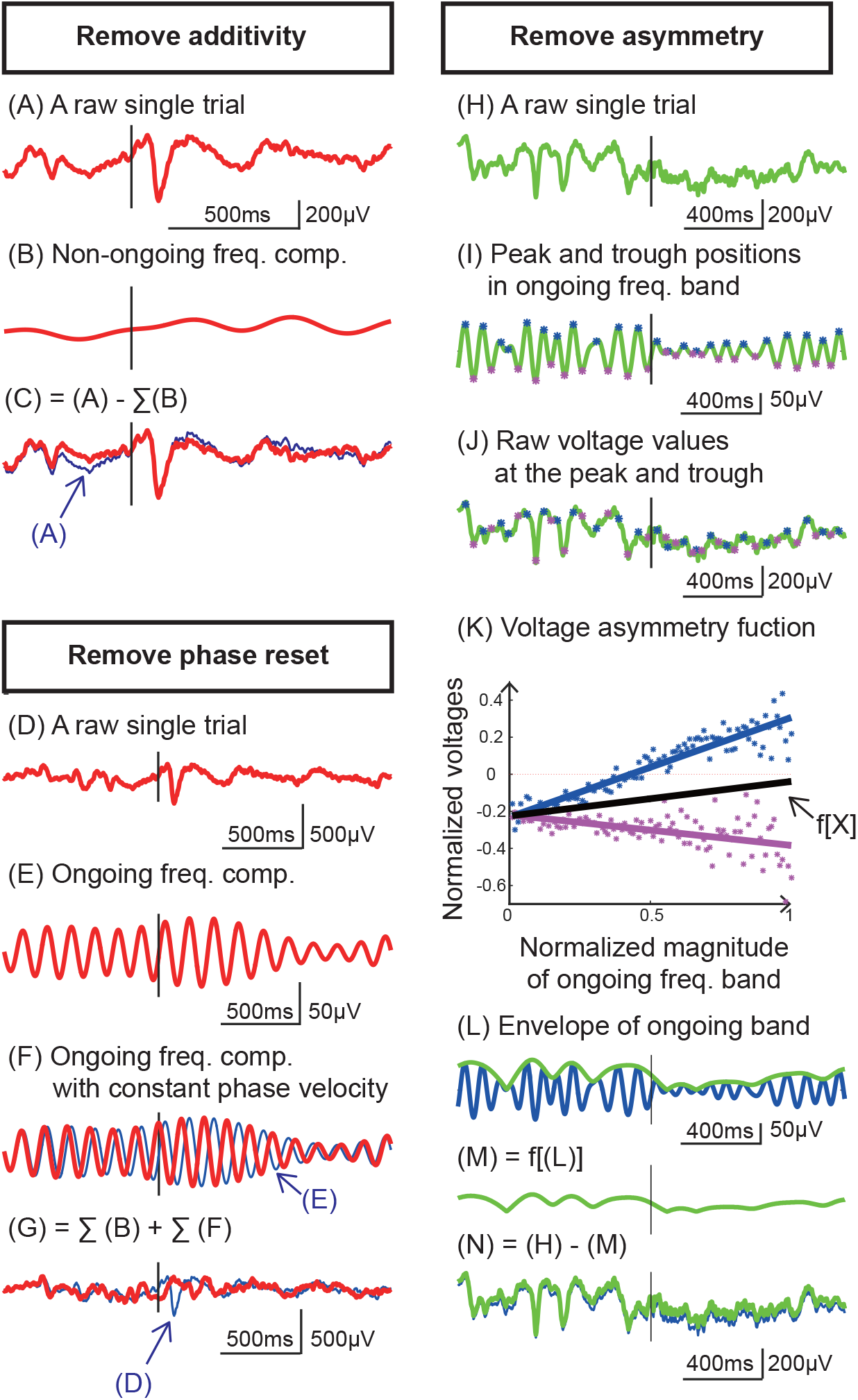
Method to remove the effect of additivity, phase reset, and asymmetric amplitude from evoked potentials. We first determined the frequency band of the ongoing oscillation (*Figure 3–Figure Supplement 1*). To remove the additive effect, we removed frequency components outside of the ongoing oscillation (see C). For this, we subtracted the frequency components outside the frequency band of the ongoing oscillation (see B) from the evoked potentials (see A). To remove the effect of phase reset, we recomposed the evoked potentials with constant phase velocity. For this, we decomposed each trial into 1 Hz-wide frequency bands between 1 and 200 Hz (see E). Next, we adjusted the signal’s ongoing oscillation to have constant phase velocity (see F). Finally, we combined the time series across all frequency bands into our recomposed signal (see G). To remove the effect of asymmetric amplitude, we subtracted the asymmetric bias from each evoked potential within the frequency band of the ongoing oscillation. For this, we first detected the peak and troughs within the ongoing oscillation (see I). Next, we determined the relationship between the amplitude at these peaks and troughs in the ongoing oscillation, and the voltage at the same time points in the original signal (see J). This analysis yielded two linear relationships (i.e., voltage-to-voltage functions), one for the peak (see the blue line in K), and one for the trough (see the purple line in K). The average between these two relationships represents the asymmetry between peak and trough amplitude as a function of the amplitude of the ongoing oscillation (see black line in K). We used this function to translate the envelope of ongoing oscillation (see L) into the asymmetric bias (see M). Finally, we subtracted the time-varying asymmetric amplitude from each trial’s original signal (see N).

In our analyses, we were interested in determining the spectral composition of the elicited AEPs and MRPs, and in quantifying the contribution of each of the three generating mechanisms to the ERP’s total energy.

The results of our spectral composition analysis shows that MRPs are comprised mostly of low frequency components (<3 Hz), while P1 and N1 of AEPs are comprised mainly of 3–40 Hz components (***Figure 3***E). Further, we observed that higher pre-stimulus oscillation power yields a bigger N1 peak amplitude in the AEP (***Figure 3***F), but doesn’t influence the MRP amplitudes (***Figure 3***G). The analysis of the specific contribution of each mechanism in the rise of the AEPs and MRPs shows that the additivity mechanism explains 61% and 88% of the energy in the AEPs and MRPs, respectively (***Figure 5***B). The phase reset mechanism explains 41% and 12% of the energy in the AEPs and MRPs, respectively (***Figure 5***B). In contrast, the asymmetric amplitude effect only explains 6% of the energy in the AEPs and MRPs (***Figure 5–Figure Supplement 1***). It should be noted that, due to the finite precision of our computations involving the removal of the three generating mechanisms from thousands of single trials, and rounding of the percentage numbers, reported aggregated energy may exceed 100%.

**Figure 5.**
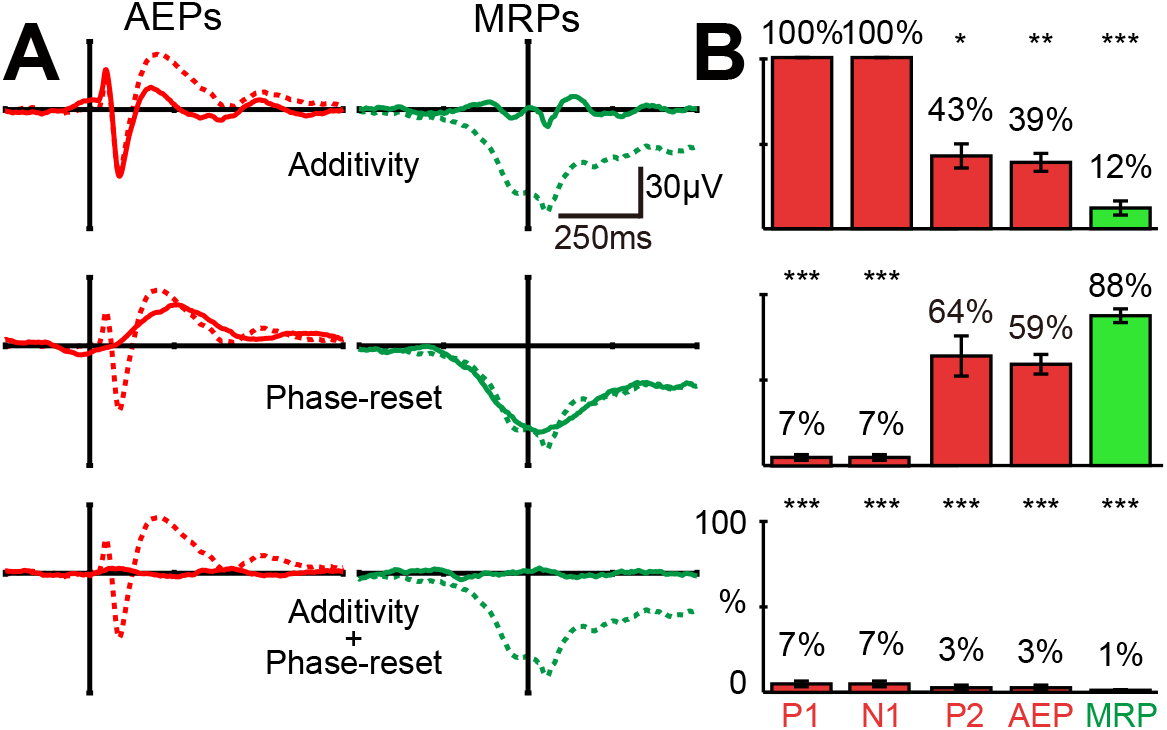
Shape (A) and energy (B) of ERP components before (dashed lines) and after (solid lines) removing additive and phase reset contribution. Additivity accounts for most of the P2 component’s energy in the AEPs (p<0.05, t-test) and almost the entire energy of the slow negative potential in the MRPs (p<0.001, t-test). Phase reset accounts for almost all of the P1 and N1 components’ energy in the AEPs (p<0.001, t-test). Together, additivity and phase reset account for almost the entire energy of AEPs and MRPs and their components (p<0.001, t-test). **Figure 5–Figure supplement 1.** Effect of power variations and asymmetric amplitude. **Figure 5–Figure supplement 2.** Asymmetric amplitude contribution to the energy of ERPs.

Finally, we found that the combination of additivity and phase reset mechanisms explains almost the entire amount of energy in the ERPs (97% and 99% of the energy in the AEPs and MRPs) (***Figure 5***). Our results show that the individual components of the AEPs (i.e., P1 and N1 components) are mostly generated by phase reset (93%). In contrast, the P2 component is generated by both additivity (57%) and phase reset (36%) (***Figure 5***B).

## Discussion

In this study, we investigated the specific contributions of additivity, phase reset, and asymmetric amplitude to AEPs and MRPs. Our results demonstrate that MRPs are generated mainly through additivity (88%) with little contribution from phase reset (12%). Within the AEPs, the P1 and N1 components are mostly generated by phase reset (93%) while the P2 component is generated by additivity (57%) and phase reset (36%). In contrast to recent speculations (***Schalk, 2015***; ***Nikulin et al., 2007***; ***Mazaheri and Jensen, 2008***), oscillatory voltage asymmetry only marginally contributes to these ERP components (6%).

### Ongoing oscillation affects ERPs and behavior

Our results show that the power of the ongoing oscillation (expressed as the pre-stimulus power in the 3–40 Hz band) directly modulates the N1 amplitude in the AEP. In contrast, the amplitude of the MRP remains unaffected by the power of low-frequency oscillations (expressed by pre-stimulus power in the <3 Hz band). This is consistent with previous studies showing that ongoing oscillations affect the AEPs (***Rahn and Basar, 1993***; ***Haig and Gordon, 1998***) and visual evoked potentials (***Makeig et al., 2002***, ***2004***; ***Min et al., 2007***). As ongoing oscillations are a hallmark of cortical inhibition, we were interested in determining whether the relationship between the amplitude of ongoing oscillations and the amplitude of the AEP’s components also extended to the resulting behavior (i.e., the reaction time to the auditory stimulus). In fact, our results show that higher power of ongoing oscillations not only increase the amplitude of the AEP’s components, but also increase reaction time (see ***Figure 3–Figure Supplement 2***). This confirms that ongoing oscillations directly affect behavior (***Klimesch et al., 2007***; ***Haegens et al., 2011***).

### Phase reset and ERPs

Our results also show that MRPs meet none of the preconditions for phase reset. First, we did not find any ongoing low-frequency oscillation (<3 Hz) within the 1000 ms-long pre-stimulus period (***Figure 3–Figure Supplement 1***). Second, the frequency characteristics of the pre-stimulus period and the MRP are different (***Figure 3***E). Third, oscillatory voltage asymmetry only marginally contributed to the generation of MRPs (***Figure 5–Figure Supplement 1***). Together with the results of our main analysis, this shows that MRPs are generated by additivity and not by phase reset (***Figure 5***).

In contrast to MRPs, our results show that AEPs meet all preconditions for phase reset. This confirms previous studies (***Makeig et al., 2002***; ***Rahn and Basar, 1993***; ***Haig and Gordon, 1998***; ***Hanslmayr et al., 2006***; ***Mäkinen et al., 2005***) that showed that the ongoing oscillation directly affects the amplitude of the AEP’s components (***Figure 3***F) and shares the same frequency characteristics (***Figure 3***E). Together with the results of our main analysis, this shows that the P1 and N1 components of the AEP are generated by phase reset and not by additivity, while the P2 component is generated jointly by additivity and phase reset (***Figure 5***). The latter result may be due to the variance in amplitude and time of the P2 component across subjects and will require further investigation (***Figure 5–Figure Supplement 2***).

### Asymmetry in oscillatory activity and ERPs

Interestingly, we did not find a significant contribution of oscillatory voltage asymmetry to the generation of ERPs (***Figure 5–Figure Supplement 1***).

On average, our ERPs were correlated with the envelope of the ongoing oscillation (Pearson’s correlation, AEPs: r^2^=0.34±0.26, MRPs: r^2^=0.44±0.23), and 18/22 auditory locations and 13/15 motor locations have significant correlations (p<0.01, bonferroni corrected). This correlation is considered to be the governing factor of the contribution of asymmetric amplitude to the generation of ERPs (***Nikulin et al., 2007***; ***Mazaheri and Jensen, 2008***; ***van Dijk et al., 2010***). An analysis of individual cortical locations confirms this, showing that this correlation is indeed a strong governing factor of the contribution of oscillatory voltage asymmetry to the generation of ERPs (Pearson’s correlation, AEPs: r=0.71, p<0.01, MRPs: r=0.80, p<0.01, as shown in ***Figure 5–Figure Supplement 2***A).

However, the shape of ERPs and the envelope of ongoing oscillation were not completely same, although there were significant correlations (see ***Figure 5–Figure Supplement 2***B). The mismatch of the ERP and the envelope of ongoing oscillation in terms of shape yields less reduction from asymmetry removal in single trials. Thus, our results confirm that oscillatory voltage asymmetry is not a prerequisite for the generation of evoked responses (***Nikulin et al., 2010***).

### Physiological interpretation

In their seminal review of the change in the ongoing EEG/MEG in the form of event-related desynchronization (ERD) or event-related synchronization (ERS), Pfurtscheller and da Silva showed that stimulus-evoked and cortex-induced activity within the cortex differ in their physiological pathways. (***Pfurtscheller and Da Silva, 1999***)

Although we only investigated auditory and motor responses in the present study, our results, along with supporting evidence from the literature, further elucidate the role of these physiological pathways. Specifically, our results in ***Figure 3***F and ***Figure 5***, along with supporting literature, (***Makeig et al., 2002***; ***Rahn and Basar, 1993***; ***Haig and Gordon, 1998***; ***Hanslmayr et al., 2006***; ***Mäkinen et al., 2005***) show that phase reset generates the P1 and N1 components within the ERPs. As P1 and N1 components are considered exogenous activity, (***Pfurtscheller and Da Silva, 1999***; ***Sur and Sinha, 2009***) phase reset can also be considered to be exogenous, or stimulus-evoked. In contrast, our results (see ***Figure 3***G and ***Figure 5***) show that additivity generates the MRP. As MRP are considered endogenous activity, (***Pfurtscheller and Da Silva, 1999***; ***Sur and Sinha, 2009***; ***Neshige et al., 1988***) additivity can also be considered endogenous (i.e., induced by cortical neurons in the absence of external stimuli). In conclusion, we infer that phase reset and additivity are involved in exogenous and endogenous activity, respectively.

### Relevance for neuronal dysfunction

The N1 response to auditory stimuli is a well-known biomarker of schizophrenia (***Ford et al., 1994***, ***2001***; ***Roth et al., 1980***; ***Javitt and Sweet, 2015***; ***Abeles and Gomez-Ramirez, 2014***). Studies investigating this biomarker also found that human subjects affected by schizophrenia, when compared to healthy subjects, not only exhibit smaller auditory N1 amplitudes, but also smaller ongoing alpha oscillations (***Abeles and Gomez-Ramirez, 2014***). Further studies found that administering Ketamine (an NDMA receptor agonist that has shown to reduce alpha power during resting state and N1 amplitudes (***de Pesters et al., 2016***; ***Javitt and Sweet, 2015***)) to healthy human subjects resulted in schizophrenia like symptoms (***Javitt, 2012***; ***Javitt and Sweet, 2015***). This suggests that alpha oscillations and N1 components share the same neural generator. Our results support this hypothesis by explaining how phase reset during smaller ongoing oscillations yields smaller N1 responses.

While our results showed that the reaction time increases with the amplitude of the ongoing oscillation ***Figure 3–Figure Supplement 2***, other studies showed that subjects affected by schizophrenia exhibit increased reaction time despite a comparatively small ongoing alpha oscillation (***Nuechterlein, 1977***; ***Kaiser et al., 2008***). These contradictory results may be explained by preoccupation of cortical resources with processing auditory hallucinations. It is well-known that a reduction in the available cortical resources directly results in a smaller observable amplitude of the ongoing oscillation (***Hanslmayr et al., 2011***; ***Pfurtscheller and Da Silva, 1999***; ***de Pesters et al., 2016***). This new insight may lead to further studies that investigate the role of ongoing oscillations in people affected by schizophrenia.

MRP and beta power are well-known biomarkers of Parkinson’s disease. Studies investigating these biomarkers have shown that subject affected by Parkinson’s disease, when compared to healthy subjects, exhibit higher beta power (***De Hemptinne et al., 2015***) and less-steeply sloped MRPs (***Cunnington et al., 1997***). Our results expand on these studies by suggesting that less-steeply sloped MRPs are a result of reduced endogenous additive activity from motor cortex due to the inhibitory effect of beta oscillations originating from sub-cortical structures.

### Limitations and potential confounds

While our experimental design controlled for many crucial confounds (i.e., through the use of precise stimulus onset and randomization of the inter-stimulus interval), several factors remained outside of our control. First, behavioral confounds, such as distractions and variations in attention, are always a possibility. However, as we recorded up to 500 trials, it is unlikely that occasional distractions or lack of attention created a systematic confound. Second, due to experimental constraints, we did not control for background noise. For that reason, we used a salient 1 kHz tone that exceeded the background noise level as our auditory stimulus. However, the saliency of the stimulus may have affected both the shape and generating mechanisms of the AEPs. Third, the subjects in this study were patients affected by epilepsy. Compared to healthy people, they may display pathological differences in responses. However, the etiology of epilepsy varied across all eight subjects, which makes a systematic confound from this source unlikely as well. Nevertheless, we took three measures to minimize this effect. First, we verified that the seizure onset zone was distant from the areas of interest. Second, we conducted the experiment when the patient was alert and had not had a seizure for at least one day. Third, we excluded cortical locations and trials that exhibited clear epileptogenic activity and verified that epileptogenic activity was not related to, or entrained by, our external auditory stimuli.

To rule out the possibility that variability in the power of the ongoing oscillation had a confounding additive effect to the generation of ERPs, we performed a control analysis in which we removed this variability from our signals. The results of this control analysis show that power variations in the ongoing oscillation can only explain 12% and 1% of the energy in the AEPs and MRPs, respectively (***Figure 5–Figure Supplement 1***).

To confirm that ERPs obtained from ECoG represented valid AEPs and MRPs, we performed a control experiment with 7 subjects in which we recorded EEG and eye-movement data. The satisfactory comparison between AEPs and MRPs obtained from ECoG and those obtained from EEG (***Figure 2–Figure Supplement 4***), as well as those described in the literature (***Figure 2–Figure Supplement 5***), allowed us to reject this potential confound. Finally, because pre-stimulus saccades are known to induce phase reset into cortical signals (***Ito et al., 2011***), we verified that the subjects maintained eye-gaze throughout the pre-stimulus period. We detected saccades from 1 s before to 1 s after stimulus onset. However, our analysis showed that only 13 of 2370 trials over 7 subjects (EEG data) exhibited saccades before stimulus onset, making this an unlikely confound.

### Future studies

Our results shed new light on the contributions of the three mechanisms (additivity, phase reset, and asymmetry) in the generation of ERPs, using signals recorded from the STG and M1 motor cortex. However, the role of the thalamus deserves more attention in the investigation of the generating mechanisms of ERPs. While our study used electrocorticographic electrodes placed on the surface of the brain, future studies could complement this with stereo-EEG electrodes placed within defined structures of the thalamus (e.g., the medial geniculate nucleus (MGN)). In general, the gyrate structure of the human brain implies that we can only subsample the cortex, and that some cortical areas remain outside of the reach of electrocorticographic recordings. For example, while we are able to record from the belt and parabelt areas of auditory cortex, most of primary auditory cortex remains inaccessible to electrocorticographic recordings. Recent progress in surgical techniques (***Kajikawa et al., 2015***; ***Jenison et al., 2015***) could overcome this limitation and allow us to extend our exploration into primary auditory regions, such as A1 primary auditory cortex. Added to our current findings, these future studies could greatly improve our understanding of the cortical and subcortical pathways involved in the generation of AEPs and MRPs (***Orrison, 2008***; ***Crosson, 1992***; ***Paradiso et al., 2004***).

## Conclusions

In summary, our results shed light on the previously unknown physiological origins of additivity, phase reset, and asymmetric amplitude mechanisms and their contribution to the generation of AEPs and MRPs. This new insight should facilitate the physiological interpretation of AEPs and MRPs. It should also guide the future creation of general models of evoked potentials and their relationship to behavior.

## Methods and Materials

### Subjects

Eight human subjects (S1–S8, 4 males, 4 females, average age = 41±14) participated in this study at the Albany Medical Center in Albany, New York. The subjects were mentally and physically capable of participating in our study (average IQ = 96±18, range 75–120, ***Wechsler 1997***). All subjects were patients with intractable epilepsy who underwent temporary placement of subdural electrode arrays to localize seizure foci prior to surgical resection.

The implanted electrode grids were approved for human use (Ad-Tech Medical Corp., Racine, WI; and PMT Corp., Chanhassen, MN). The platinum-iridium electrodes were 4 mm in diameter (2.3 mm exposed), spaced 10 mm center-to-center, and embedded in silicone. The electrode grids were implanted in the left hemisphere for seven subjects (S1, S3, S6, and S7) and the right hemisphere for five subjects (S2, S4, S5 and S8). Following the placement of the subdural grids, each subject had postoperative anterior-posterior and lateral radiographs, as well as computer tomography (CT) scans, to verify grid location. These CT images, in conjunction with magnetic resonance imaging (MRI), were used to construct three-dimensional subject-specific cortical models and derive the electrode locations (***Coon et al., 2016***).

Patients with ECoG coverage extending from auditory to motor cortex are a relatively rare occurrence. Thus, most previous ECoG studies were limited to subjects with either motor or auditory coverage and typically reported results from less than 8 participants (***Jenison et al., 2015***; ***Edwards et al., 2005***; ***Paradiso et al., 2004***; ***Neshige et al., 1988***). In our study, we recorded electrocorticographic signals from auditory and motor cortex in 8 human subjects (***Figure 2–Figure Supplement 1***).

A further seven human subjects (X1–X7, all males, average age = 27±3.6) served as a control group for which we recorded EEG and eye-movement data. These subjects were fitted with an elastic cap (Electro-cap International, ***Blom and Anneveldt 1982***) with tin (***Polich and Lawson, 1985***) scalp electrodes in 64 positions according to the modified 10-20 system (***Sharbrough, 1991***).

All subjects provided informed consent for participating in the study, which was approved by the Institutional Review Board of Albany Medical College and the Human Research Protections Offce of the U.S. Army Medical Research and Materiel Command.

### Data collection

We recorded ECoG signals from the subjects at their bedside using the general purpose Brain-Computer Interface (BCI2000) software (***Schalk et al., 2004***), interfaced with eight 16-channel g.USBamp biosignal acquisition devices, or one 256-channel g.HIamp biosignal acquisition device (g.tec., Graz, Austria) to amplify, digitize (sampling rate 1,200 Hz) and store the signals. To ensure safe clinical monitoring during the experimental tasks, a connector split the cables connected to the patients into a subset connected to the clinical monitoring system and a subset connected to the amplifiers.

We recorded EEG signals and eye-movement coordinates from the subjects in our control group using the same g.USBamp setup and a Tobii T60 eye-tracking monitor (Tobii Tech., Stockholm, Sweden) that was positioned at eye level 55–60 cm in front of the subject and was calibrated for each subject at the start of each experimental session.

### Task

The subjects performed an auditory reaction task in which they responded with a button press to a salient 1 kHz tone. For this, the subjects used their thumb contralateral to their ECoG implant. In total, the subjects performed between 134 and 580 trials. Throughout each trial, the subjects were first required to fixate gaze onto the screen in front of them. Next, a visual cue indicated the start of the trial, which was followed by a random 1–3 s pre-stimulus interval and subsequently, the auditory stimulus. The stimulus was terminated by the subject’s button press, or after a 2 s time out, after which the subject received feedback about his/her reaction time. This feedback motivated the subjects to respond as fast as possible to the stimulus. To prevent false starts, we penalized subjects with a warning tone if they responded too fast (i.e., less than 100 ms after stimulus onset). We excluded false-start trials from our analysis. In this study, we were interested in the auditory and motor response to this task. This required defining the onset of these two responses. For the auditory response, we defined this as the onset of the auditory stimulus (as measured by the voltage between the sound port on the PC and the loudspeaker). For the motor response, we defined the onset as the time when the push-button was pressed. To ensure the temporal accuracy of these two onset markers, we sampled them simultaneously with the ECoG signals using dedicated inputs in our biosignal acquisition system. We defined baseline and task periods for the auditory and motor response. Specifically, we used the 0.5-s period prior to the stimulus onset as the baseline for the auditory response, and the 1-s to 0.5-s period prior to the button press as the baseline for the motor response. Similarly, we used the 1-s period after stimulus onset as the task period for the auditory response, and the period from 0.5-s before to 0.5-s after the button press as the task period for the motor task.

### Data pre-processing

As our amplifiers acquired raw unfiltered ECoG signals, we first removed any offset from our signals using 2nd order Butterworth highpass filter at 0.05 Hz. Next, we removed any common noise, using a common median reference filter. For the creation of the common-mode reference, we excluded signals that exhibited an excessive 60 Hz line noise level (i.e., ten times the median absolute deviation). To improve the signal-to-noise ratio of our recordings and to reduce the computational complexity of our subsequent analysis, we downsampled our signals from 1200 to 400 Hz using MATLABs “resample” function, which uses a polyphase antialiasing filter to resample the signal at the uniform sample rate.

### Electrode selection

To select the appropriate electrodes, we needed to determine which electrodes exhibited an early auditory or motor-related response. We accomplished this in three steps. In the first step, we selected only those cortical locations that exhibited a task-related response in the high gamma band. For this purpose, we performed a statistical comparison between baseline and task periods across all trials. Specifically, we calculated the Spearman’s correlation coeffcient between the power of baseline/task periods, and a corresponding label (i.e., −1 for baseline and +1 for task period). This yielded one correlation value for each of our cortical locations. Next, we performed a permutation test to determine the significance of each cortical location’s correlation value. In this, we calculated our correlation coefficient 1000 times, each time with a newly permutated sequence of labels. This yielded a distribution of correlation values with an area under the curve (AUC) of 1.

Next, we determined the significance of our true correlation value as the single-tailed AUC created by the intersection of the true correlation value with the distribution of correlation values obtained from the permutation test. Finally, we identified the cortical locations exhibiting a task-related response in the high gamma band that had a p-value smaller than 0.001 (Bonferroni-corrected for the number of cortical locations). In the second step, for each subject, we restricted our selection to the single auditory and single motor-related cortical locations that exhibited the earliest onset. To perform this selection, we first calculated the Spearman’s correlation coeffcient between the power of baseline/task periods for individual time points of the task period (i.e., auditory: 500 ms after stimulus onset for auditory; motor: 250 ms before button press). Next, we determined the onset of task-related cortical activity as the earliest time point when the correlation exceeded the 99th percentile of all correlation values. This defined one auditory-related and one motor-related location exhibiting the earliest cortical onset. Finally, in the third step, we expanded our selection to include those cortical locations, identified in the first step, that were within 15 mm of our earliest onset locations.

### Removing effects of phase reset

Phase reset is reflected as spikes in the phase velocity, i.e., the first derivative of the phase signal (***Freeman et al., 2003***, ***2006***; ***Thatcher, 2012***). To remove the effect of phase reset, we recomposed each trial’s original signal with constant phase velocity (see ***Figure 4***). To accomplish this, we first decomposed each trial’s signal into 1 Hz wide frequency bands between 1 and 200 Hz. For this, we first applied a fast Fourier transform (FFT) on each trial’s signal to calculate the discrete Fourier transform (DFT) of our signals. This decomposition yielded the frequency representation of our signal. We then used the inverse FFT to recreate individual time series for each 1 Hz wide frequency band. Next, we determined the constant velocity that each frequency band’s time series should have as its phase velocity during the baseline period (calculated as a first derivative of the phase of the Hilbert transform, as shown in ***Figure 4***E). Next, for frequency bands above 3 Hz, we recomposed the time series with constant phase velocity. For this, we applied the envelope from the original time series onto a cosine signal with the constant phase velocity determined in the previous step. Finally, we applied an inverse FFT on each frequency band to recompose the signal. This recomposition yielded a time series with constant phase and the same power as the original time series. Finally, we combined the time series across all frequency bands into our recomposed signal.

### Removing effects of asymmetric amplitude

Asymmetric amplitude is characterized by an asymmetry between the peak and trough amplitudes of ongoing oscillations. This asymmetry varies over time and can affect the shape of low-frequency components within the ERP (***Mazaheri and Jensen, 2008***). To remove this effect, we estimated and removed the time-varying asymmetric amplitude from each trial’s original signal (see ***Figure 4***H–N). As the asymmetry between peak and trough amplitude is a function of the peak and trough amplitudes themselves (***Mazaheri and Jensen, 2008***), we first needed to determine this function for each cortical location. For this purpose, we first detected the peak and troughs in the ongoing oscillation (i.e., the 3–40 Hz filtered signal). Next, we identified the relationship between the amplitude at these peaks and troughs in the ongoing oscillation and the voltage at the same time points in the original signal. This analysis yielded two linear relationships (i.e., voltage-to-voltage functions), one for the peak, and one for the trough. The average between these two relationships represents the asymmetry between peak and trough amplitude as a function of the amplitude of the ongoing oscillation. We use this function to determine and subtract the time-varying asymmetric amplitude from each trial’s original signal.

### Removing effect of variability in signal power

Variability in the power of the ongoing oscillation can affect the shape of the resulting ERP (***Makeig et al., 2002***; ***Min et al., 2007***). To account for this potentially confounding effect, we removed this variability from our signals. To accomplish this, we recomposed each trial’s original signal with constant power, as described in the previous section on removing the effects of phase reset. However, for this recomposition, we applied constant amplitude instead of constant phase. We determined the constant amplitude that each frequency band’s time series should have, as its median amplitude during the baseline period. This recomposition yielded a time series with constant amplitude and the same phase as the original time series. We combined the time series across all frequency bands into our recomposed signal.

## Data and Code Availability

The dataset and code accompanying this manuscript have been deposited in an online repository (https://doi.org/10.5281/zenodo.4361654). Access will be granted by the Corresponding Author upon reasonable request, or without restrictions after the publication of this manuscript.

## Acknowledgments

The authors thank Drs. Scott Makeig and Ole Jensen for their invaluable feedback. This work was supported by the NIH/NIBIB (P41-EB018783, R01-EB026439), the NIH/NINDS (U01-NS108916 and U24-NS109103), the NIH/NIMH (P50-MH109429), the US Army Research Office (W911NF-07-1-0415, W911NF-08-1-0216 and W911NF-14-1-0440), Fondazione Neurone, and the Institute of Information & Communications Technology Planning & Evaluation (IITP) grant funded by the Korea government (No. 2017-0-00451).

## Author contributions

Conceptualization: H.C., G.S. and P.B.; Methodology: H.C., G.S., L.M., S.C.J. and P.B.; Software: M.A., W.G.C. and P.B.; Validation: P.B.; Formal Analysis: H.C.; Investigation: H.C., M.A., L.M., W.G.C. and P.B.; Resources: P.B.; Data Curation: P.B.; Writing-Original Draft: P.B.; Writing-Review and Editing: H.C., M.A., J.R.W. and P.B.; Visualization: H.C. and M.A.; Supervision: G.S., S.C.J. and P.B.; Project Administration: G.S., J.R.W. and P.B.; Funding Acquisition: G.S., S.C.J., J.R.W. and P.B.;

## Appendix 1

**Appendix 1 Figure 1.**
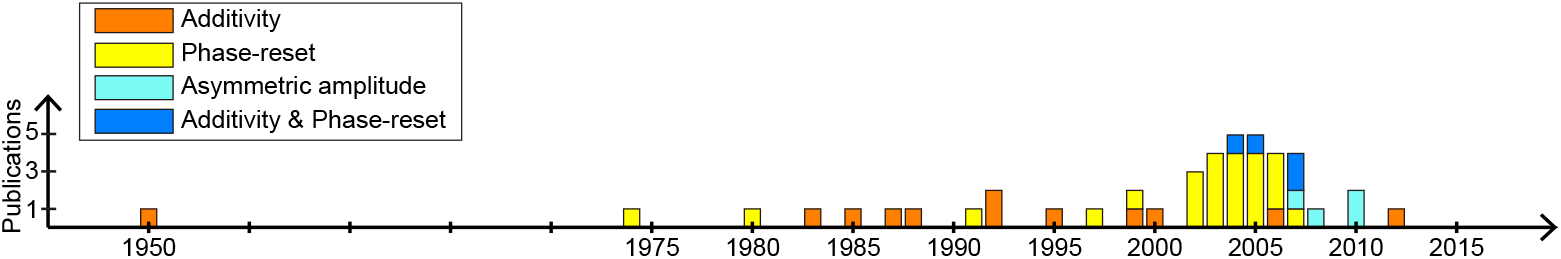
Number of scientific studies investigating the role of additivity, phase reset and asymmetric amplitude in the generation of ERPs. The controversy regarding the roles of additivity and phase reset in the generation of ERPs has been ongoing throughout the 20th century. Seminal work by Makeig et al. in 2002 (***Makeig et al., 2002***) sparked a series of combined studies which concluded that both additivity and phase reset play an important role in the generation of ERPs. In addition, seminal work by Nikulin et al. (***Nikulin et al., 2007***) culminated in the confirmation of asymmetric amplitude as a third generating mechanism of ERPs.

---

**Figure 1–Figure supplement 1.**
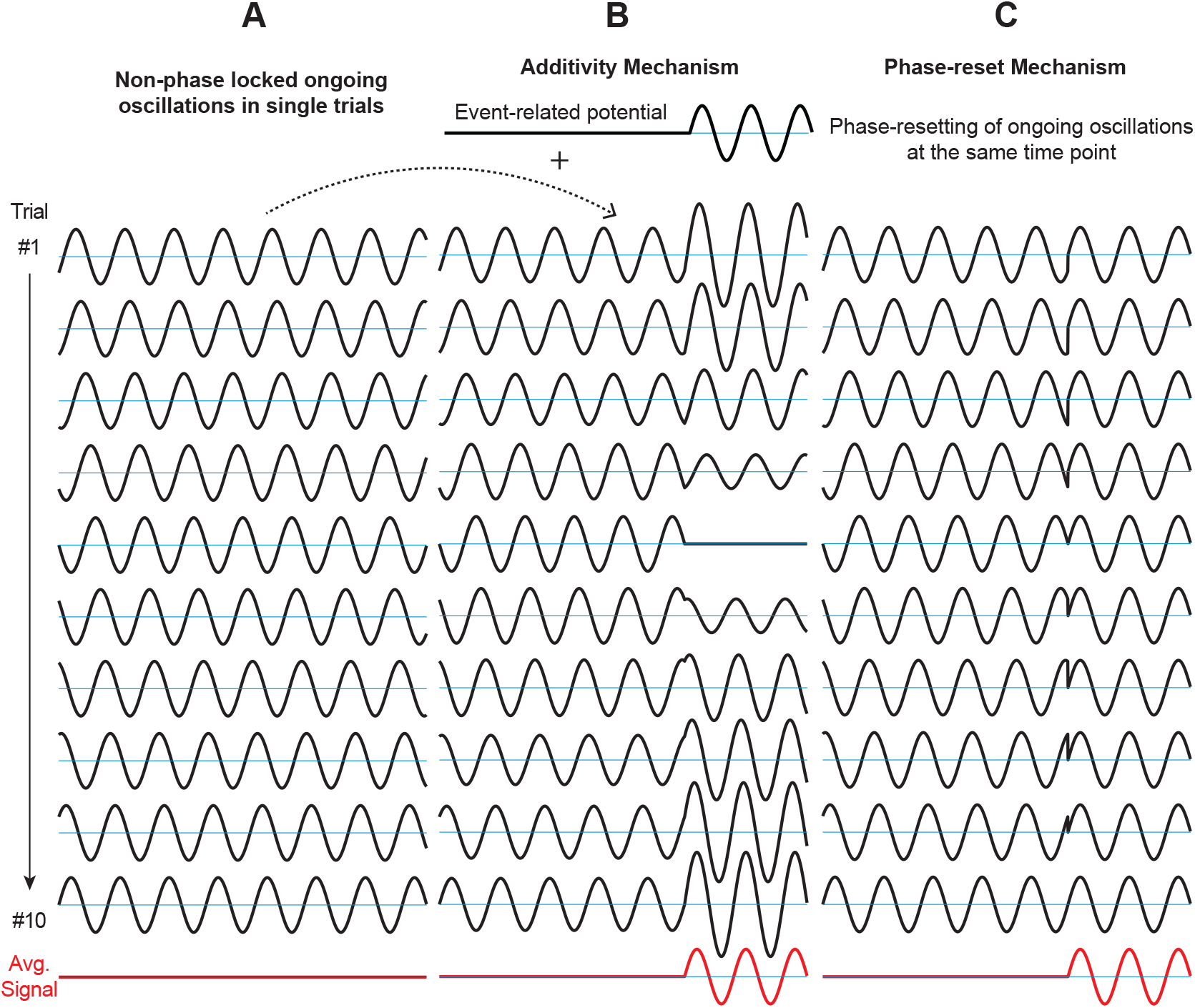
Graphical explanation of the role of additivity and phase reset in generating event-related potentials (ERPs). The oscillations depicted in A–C are non-phase-locked across trials. **A.** Without any change to the amplitude or phase of the signals, the average across trials equals zero. **B.** In the additivity mechanism, in each trial, components exhibiting the same frequency characteristic as the ongoing oscillation are added to the ongoing oscillations. The average across trials reveals an ERP. The main characteristic of additivity is the independence between ongoing oscillation and evoked potential. Thus, ongoing oscillations are considered background noise that averages out to zero across trials, and only external stimuli directly affect ERPs. **C.** In the phase-reset mechanism, for each trial, the phase of the ongoing oscillation is reset at a certain time point after stimulus onset. The average across trials reveals an ERP. In contrast to additivity, the generation of an ERP is dependent on the phase of the ongoing oscillation. Thus, external stimuli can affect ERPs only indirectly through inducing a phase-reset in the ongoing oscillation. While the additivity and phase-reset mechanisms can both explain the generation of ERPs, their physiological pathways may be fundamentally different.

**Figure 1–Figure supplement 2.**
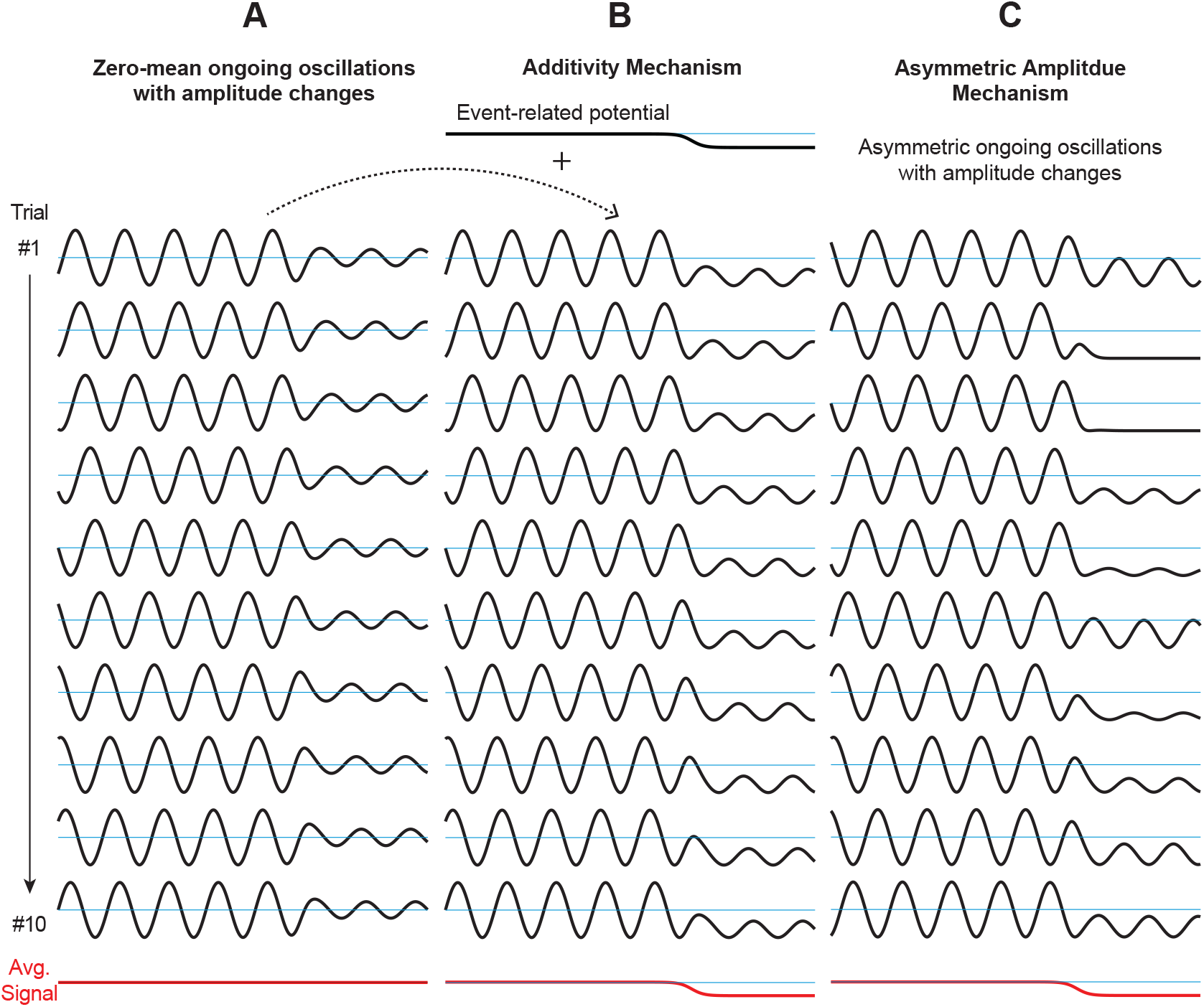
Graphical explanation of the role of additivity and asymmetry in generating event-related potentials (ERPs). The oscillations depicted in A–C are non-phase-locked across trials. **A.** Without any change to the bias of the signals, the average across trials equals zero. **B.** In the additivity mechanism, in each trial, an ERP is added to the ongoing oscillation. The average across trials reveals the ERP, while individual trials appear to exhibit asymmetry. **C.** In the asymmetric amplitude mechanism, for each trial, the ongoing oscillation exhibits asymmetry during or after the amplitude changes. The average across trials reveals the ERP. In contrast to additivity, the generation of an ERP is dependent on the amplitude envelope of the ongoing oscillation. Thus, external stimuli can affect ERPs only indirectly through affecting the amplitude and asymmetry in the ongoing oscillation. While the additivity and asymmetric amplitude mechanisms can both explain the generation of ERPs, their physiological pathways may be fundamentally different.

**Figure 2–Figure supplement 1.**
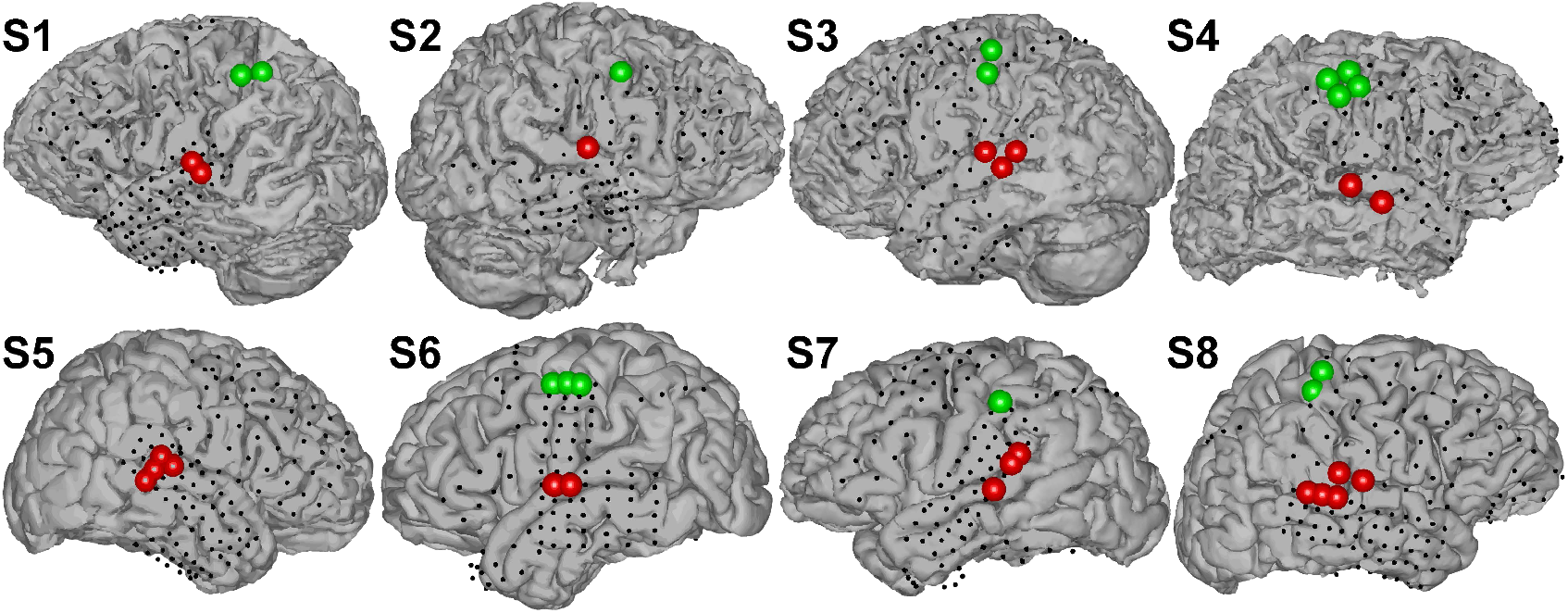
Cortical locations with activation in the high gamma band (70–170 Hz) during auditory (red) and motor task (green). Auditory tasks activated areas in STG, while motor tasks activated areas in M1 motor cortex.

**Figure 2–Figure supplement 2.**
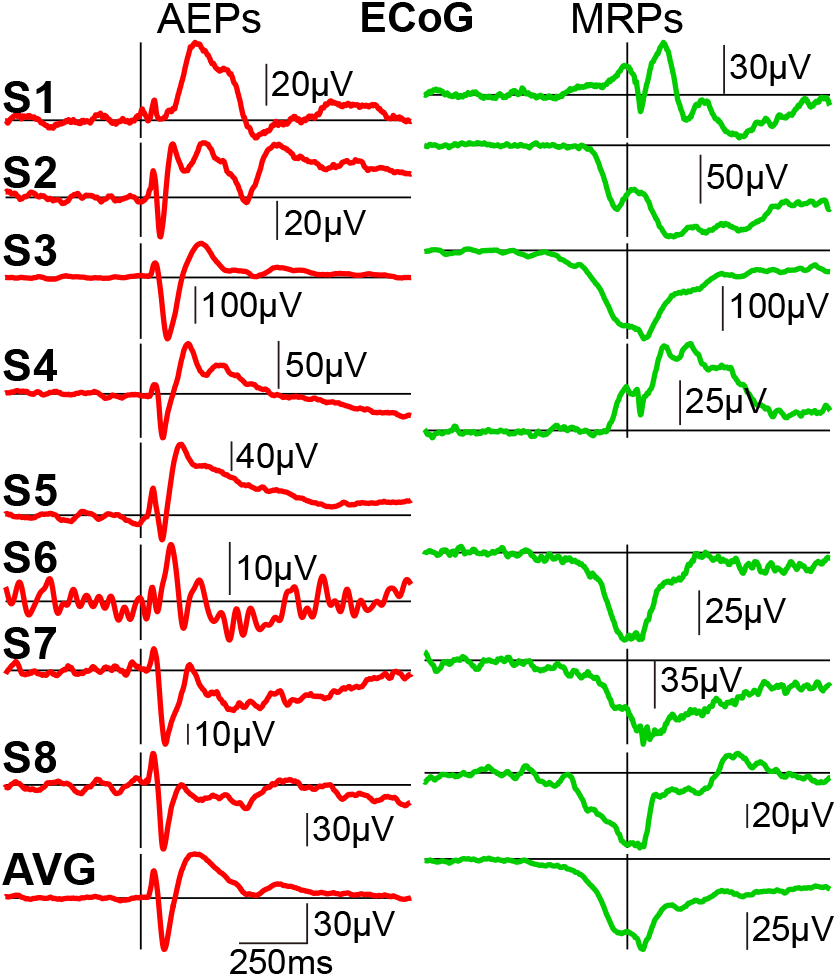
AEPs (left) and MRPs (right) from ECoG responses in subjects S1–S8. AEPs show typical N1, P1, and P2 components in 6/8 subjects. MRPs show a typical slow negative potential at movement onset in 5/7 subjects. The amplitudes of these potentials vary across subjects.

**Figure 2–Figure supplement 3.**
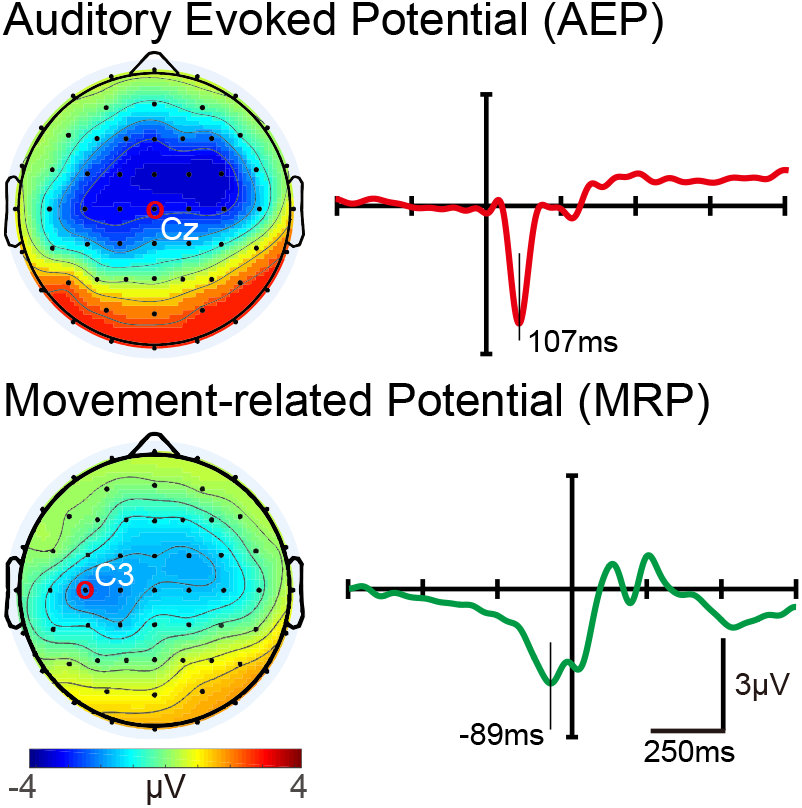
Topography and shape of grand average AEP (top) and MRP (bottom) obtained from EEG across seven subjects in this study. Topographies (left) depict the voltage distribution at the peak of the AEP (107 ms) and MRP (−89 ms), respectively. AEP and MRP plots are centered at stimulus onset and button press, respectively.

**Figure 2–Figure supplement 4.**
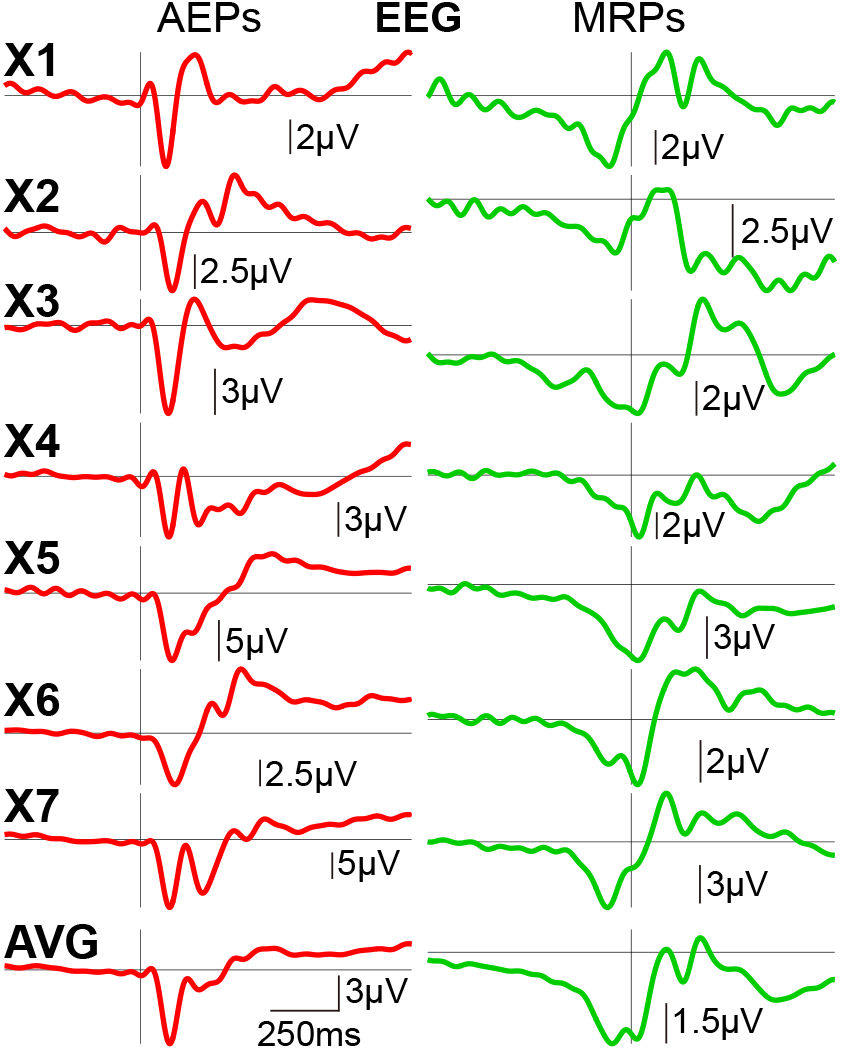
AEPs at Cz (left) and MRPs at C3 (right) from EEG responses in subjects X1-X7. AEPs show typical N1, P1, and P2 components in all subjects. MRPs show a typical slow negative potential at movement onset in all subjects. There is very little variance in the amplitude of these responses across subjects.

**Figure 2–Figure supplement 5.**
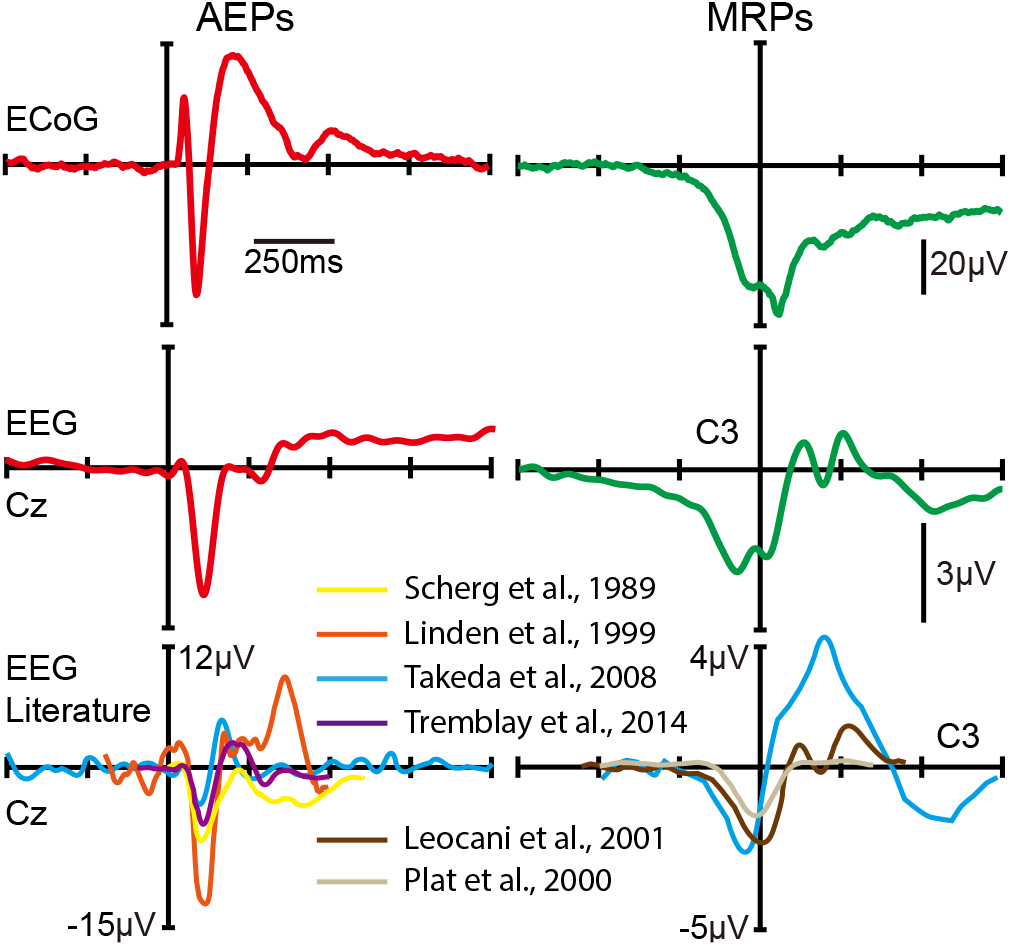
Grand average ERPs obtained from ECoG and EEG in this study (top and center) and grand average ERPs from the EEG literature (bottom). The shape of ERPs obtained from ECoG is in general agreement with ERPs obtained from EEG and with the description of ERPs in the EEG literature (***Scherg et al., 1989***; ***Linden et al., 1999***; ***Takeda et al., 2008***; ***Tremblay et al., 2014***; ***Leocani et al., 2001***; ***Plat et al., 2000***). Specifically, AEPs obtained from ECoG demonstrate the same N1, P1, and P2 components, as previously described in the EEG literature. Similarly, MRPs obtained from ECoG demonstrate the same slow negative potential around movement onset time.

**Figure 3–Figure supplement 1.**
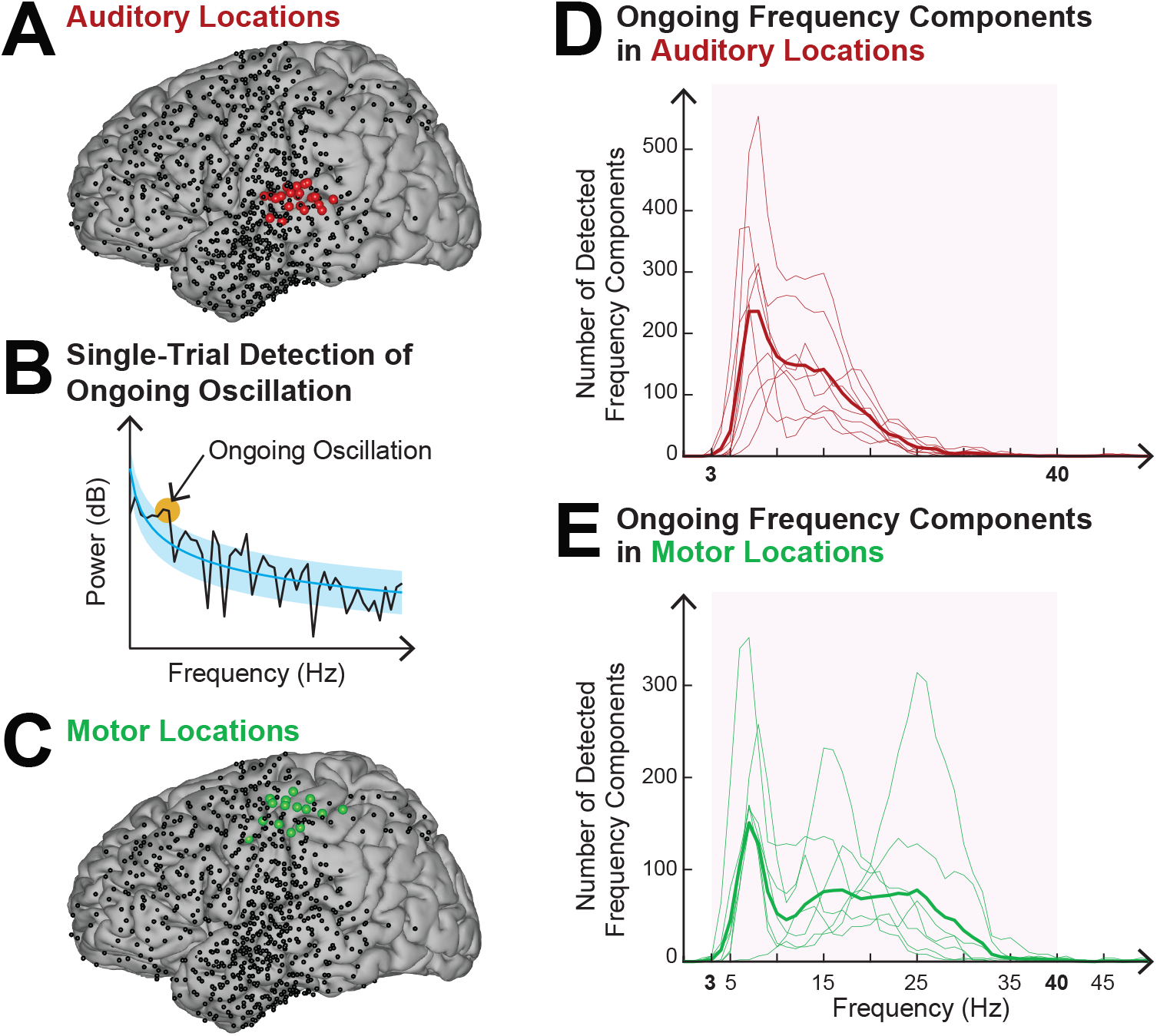
Determining the frequency band of the ongoing oscillation for auditory and motor across all eight subjects and all trials. **A.** Electrode locations for all subjects (black dots). Locations that exhibited task-related activity during auditory stimulation are highlighted in red. **B.** Detecting frequency of ongoing oscillations. Ongoing oscillations are detected as those clusters of points (orange) in the power spectrum (black) that exceed one *<Y* (blue-shaded) of the estimated 1/f power-law spectrum (blue, applied in single trials to the 1000 ms-long pre-stimulus period, see ***Donoghue et al. 2020***; ***Buzsáki et al. 2013***; ***Nunez and Srinivasan 1981*** for details). **C.** Locations that exhibited task-related activity during motor movements are highlighted in green. **D.** & **E.** Histograms of detected ongoing oscillation frequencies for auditory (D, red) and motor (E, green) locations. Thin lines represent individual subjects, and thick lines the average across subjects. The red shaded area depicts the frequency of the ongoing oscillation. Within this band, most ongoing oscillations occur at 7 Hz. Motor locations exhibit ongoing oscillations in the beta band. This analysis determined the frequency band of ongoing oscillation as 3–40 Hz.

**Figure 3–Figure supplement 2.**
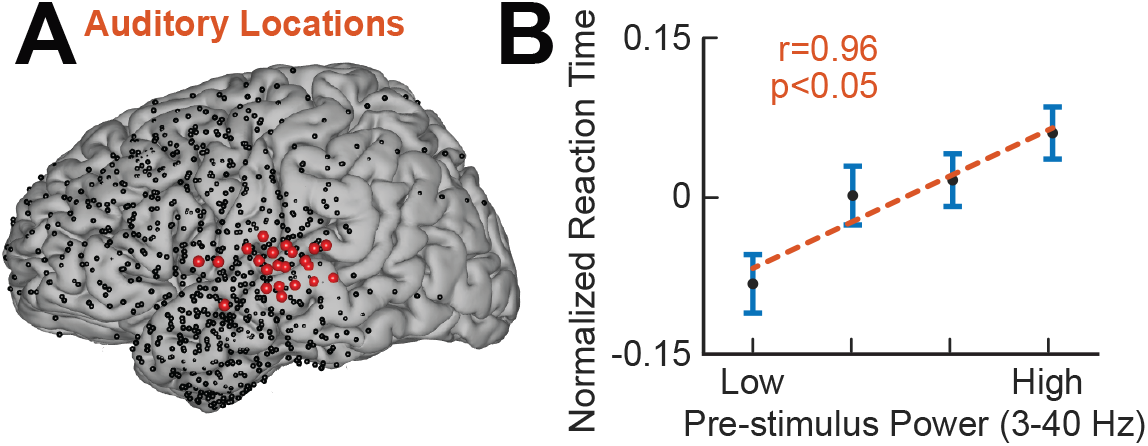
Relationship between ongoing oscillatory power and reaction time. **A.** Electrode locations for all subjects (black dots). Locations exhibiting a high gamma response (70–170 Hz) within auditory cortex are depicted in red. **B.** Increased pre-stimulus (−250–0 ms) oscillatory power (3–40 Hz) increases reaction time (r=0.96, p<0.05, Pearson’s correlation, binned pre-stimulus 3–40 Hz power, across all trials and all subjects).

**Figure 5–Figure supplement 1.**
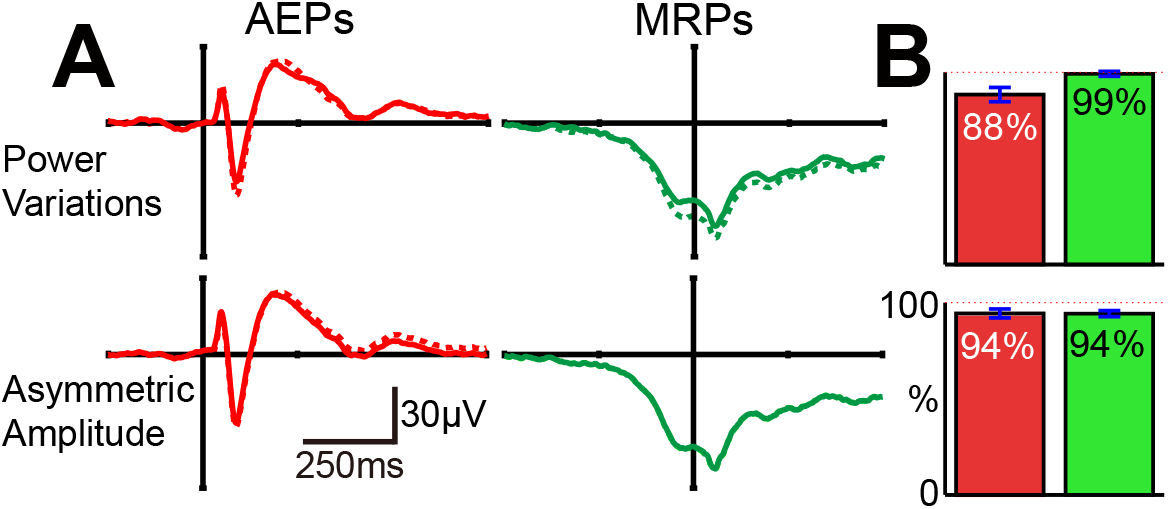
Effect of power variations and asymmetric amplitude of the ongoing oscillation on the shape and energy of ERPs. **A.** Shape of AEPs and MRPs before (dashed lines) and after (solid lines) removing power variations and asymmetric amplitude from the ongoing oscillation. Neither AEPs nor MRPs are markedly affected in their shape by this removal. **B.** Energy remaining in the resulting AEPs and MRPs after removing power variations and asymmetric amplitude from the ongoing oscillation. Neither AEPs nor MRPs are significantly affected in their energy by this removal. Removing the contribution of power variations reduces the energy of the resulting AEPs, while the energy of MRPs remains unaffected. In contrast, removing the contribution of asymmetric amplitude minimally and equally reduces the energy of AEPs and MRPs.

**Figure 5–Figure supplement 2.**
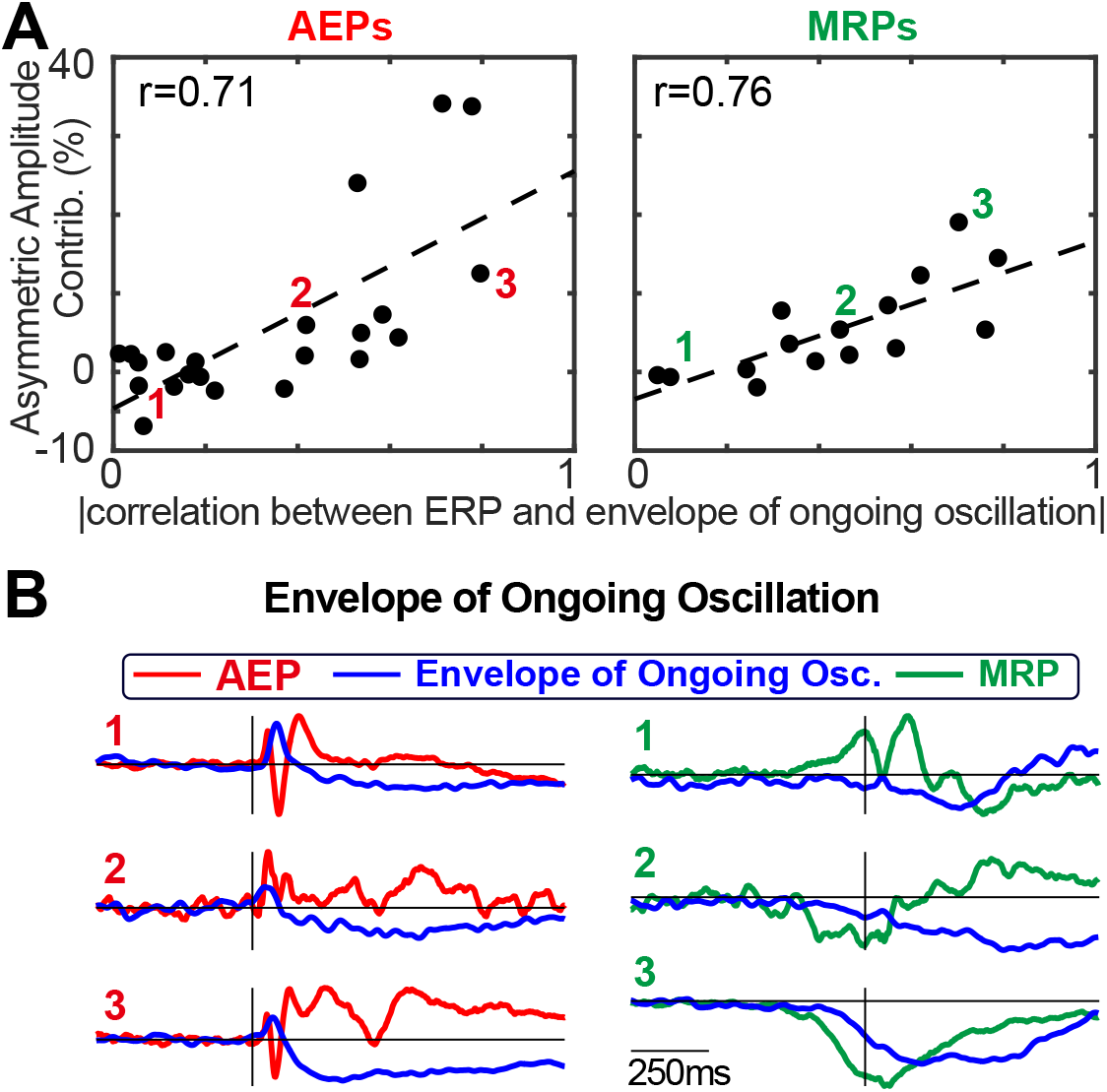
Asymmetric amplitude contribution to the energy of ERPs. **A.** The contribution of asymmetric amplitude to the energy of ERPs is driven by the ERP’s correlation with the envelope of the ongoing oscillation (AEPs: p<0.01, r=0.71; MRPs: p<0.01, r=0.76; Pearson’s correlation). **B.** Three examples for low, medium, and high correlations between ERP and the envelope of the ongoing oscillation.

